# A Universal Proximity CRISPR Cas12a Assay for Ultrasensitive Detection of Nucleic Acids and Proteins

**DOI:** 10.1101/734582

**Authors:** Yongya Li, Hayam Mansour, Yanan Tang, Feng Li

## Abstract

Herein, we describe a proximity CRISPR Cas12a assay that harnesses “collateral” single-stranded DNase activity of Cas12a as a universal amplifier for the ultrasensitive detection of nucleic acids and proteins. The target recognition is achieved through proximity binding rather than recognition by CRISPR RNA (crRNA), which allows the flexible assay design and expansion to proteins. A binding-induced primer extension reaction is then used to generate a predesigned CRISPR-targetable sequence as a barcode for signal amplification. We demonstrate that our assay is highly sensitive and universal. As low as 1 fM nucleic acid target could be detected isothermally in a homogeneous solution via the integration with nicking cleavage. We’ve also successfully adapted the assay for the sensitive and wash-free detection of antibodies in both buffer and diluted human serum samples.

## Introduction

Signal amplification is the key to the ultrasensitive detection of trace levels of biomarkers in biological and clinical samples. Recent advances in microbial clustered regularly interspaced short palindromic repeats (CRISPR) and CRISPR-associated (CRISPR-Cas) enzymes offer exciting opportunities for developing novel signal amplifiers that are both sensitive and specific.^1–5^ The RNA cleaving nuclease Cas13a was found to possess an indiscriminative ribonuclease activity upon the recognition of specific RNA targets.^2^ By combining the “collateral” cleavage of Cas13a with recombinase polymerase amplification (RPA), Zhang and co-workers introduced a series of specific high-sensitivity enzymatic reporter unlocking (SHERLOCK) techniques for the ultrasensitive detection of nucleic acids amenable for point-of-care (POC) applications.^3–5^ Doudna and coworkers have revealed the similar “collateral effect” of Cas12a, where single-stranded deoxyribonuclease (ssDNase) activities of Cas12a can be activated upon target recognition.^6^ A DNA endonuclease-targeted CRISPR trans reporter (DETECTR) system was further introduced via the integration with RPA, which enabled the detection of target DNA with attomolar sensitivity.^6^ So far, the detection of each specific genetic marker requires the design of a new CRISPR RNA (crRNA), which can be expensive and time-consuming. More importantly, it is not yet possible to expand such powerful detection systems to non-nucleic-acid targets, such as proteins. Herein, we introduce an alternative proximity CRISPR Cas12a strategy, where the target nucleic acid or protein will be translated into a universal pre-designed CRISPR-targetable DNA barcode in homogeneous solutions through a binding-induced primer extension reaction. By doing so, we are able to decouple the target recognition from the CRISPR Cas12a amplification and thus allow the broad adaptation to diverse DNA, RNA, and non-nucleic-acid targets.

## Results and Discussion

As illustrated in Scheme 1, the CRISRP-targetable protospacer-adjacent motif (PAM) containing DNA barcode for activating Cas12a is generated in situ through a binding-induced primer extension reaction between P1 and P2. P2 is coded with a pre-designed complementary sequence of the barcode. P1 and P2 contain a short 6-nt complementary sequence (red domains) with an estimated melting temperature (Tm) of 17 °C. Therefore, P1 and P2 do not hybridize under our assay condition (37 °C). However, in the presence of the target nucleic acid, P1 and P2 hybridize adjacently to the same target through complementary sequences (black domains), leading to the formation of a three-way junction with an estimated Tm of 37 °C (by NuPack). A primer extension reaction is then triggered to extend P1 to produce the DNA barcode which is recognized by the pre-designed crRNA. The ssDNase activity of Cas12a is then activated. A fluorescence turn-on assay can then be achieved by adding a short ssDNA substrate labeled with a fluorophore at the 5’ end and a quencher at the 3’ end, respectively. As the generation of the DNA barcode is quantitatively determined by the amount of the original target, the fluorescence signal can be used for target quantification.

**Scheme 1.**
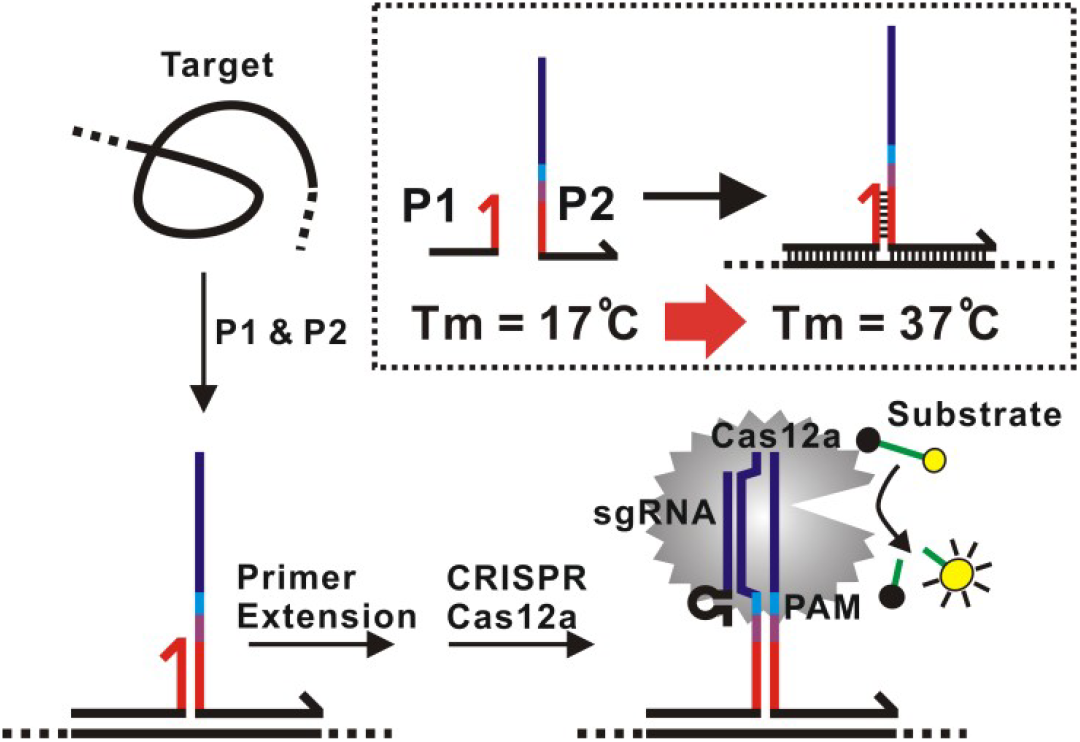
Schematic illustration of the proximity CRISPR Cas12a assay for the amplified detection of nucleic acids.

We first set out to establish the proximity CRISPR Cas12a assay for the detection of nucleic acids (Fig. 1). Upon the binding-induced primer extension followed by Cas12a cleavage, we were able to detect a synthetic target DNA with a limit of detection (LOD) at 10 pM (Fig. 1B and 1C). The LOD of our assay is comparable with that using direct crRNA recognition and Cas12a cleavage (Fig. S1), suggesting that the binding-induced primer extension can effectively translate the detection of a target sequence to the production of the DNA barcode for the subsequent Cas12a-mediated amplification. To further push the LOD of our assay, we next integrated a nicking cleavage mechanism into the production of the DNA barcode (Fig. 1A). Specifically, P2 was designed to contain a nicking recognition domain (purple) adjacent to PAM. Upon primer extension, a sequence that contains both the barcode for Cas12a activation and a nicking cleavage domain was generated. In the presence of a nicking endonuclease, the barcode was released through nicking cleavage and a new round of primer extension. As such, each target DNA could trigger the production of multiple single-stranded DNA barcodes that activate Cas12a ssDNase in a PAM-independent manner and thus further amplify the detection signal.

**Fig. 1.**
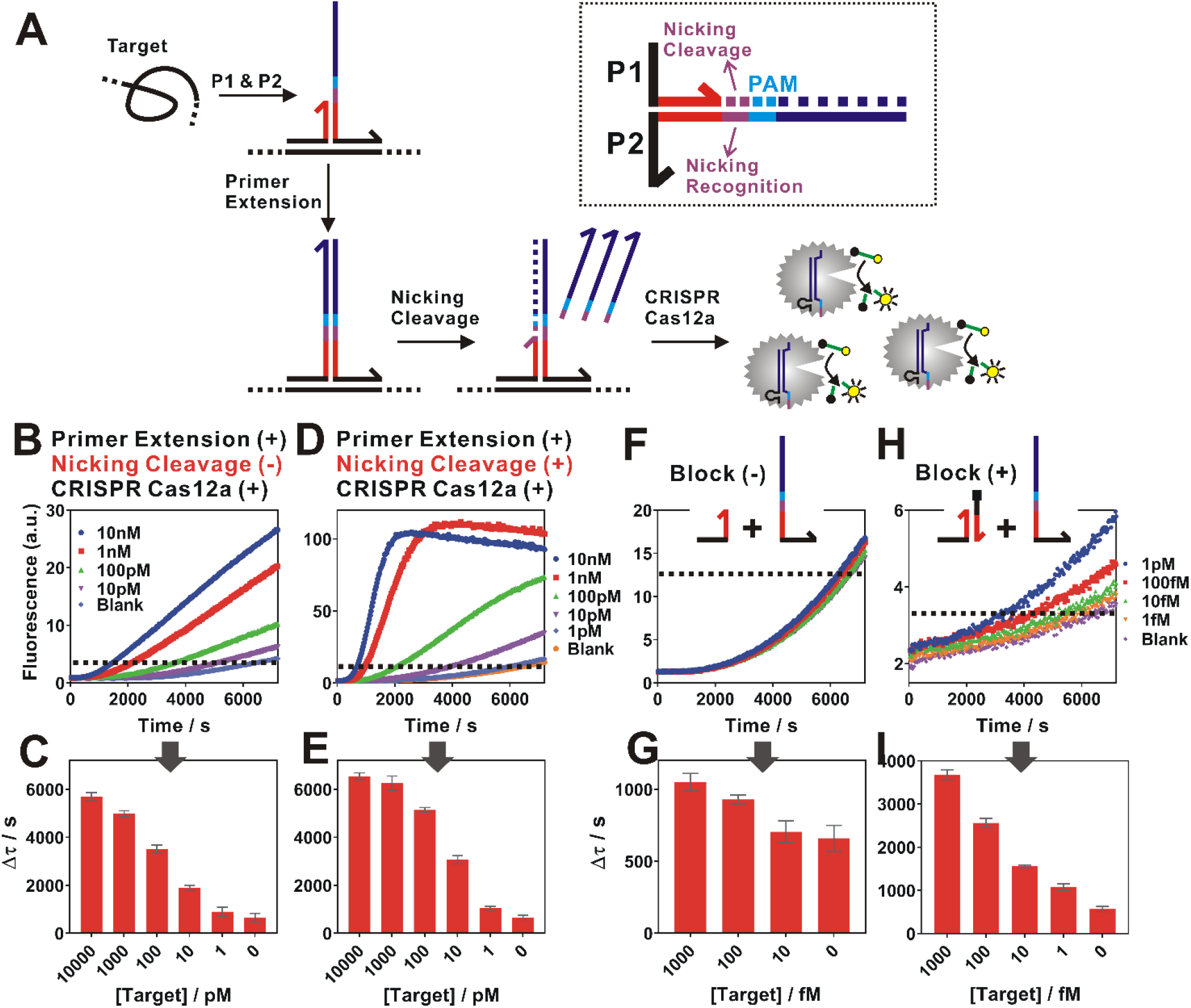
Proximity CRISPR Cas12a assay for the detection of nucleic acids. (A) Schematic illustration of the integration of proximity CRISPR Cas12a assay with nicking cleavage for further signal amplification. (B, C) Detection of a synthetic nucleic acid target using proximity primer extension followed by CRISPR Cas12a amplification. Once measuring the fluorescence increase in real-time (B), we set a threshold (dashed line) to determine the critical time τ, which is the minimal time to reach the threshold. A calibration curve was then established by plotting Δτ (Δτ = 7200s – τ) as a function of target concentrations (C). (D, E) Detection of nucleic acid target by integrating the proximity CRISPR Cas12a assay with nicking cleavage. (F, G) The detection of target at concentrations from 1 fM to 1 pM using the proximity CRISPR Cas12a assay integrated with nicking cleavage. To further push the detection limit to lower target concentrations, a blocking DNA was introduced to suppress the background (H, I). Each error bar represents one standard deviation from triplicate analyses.

As shown in Fig. 1D, a drastic enhancement in sensitivity was observed when integrating the proximity CRISPR Cas12a amplification with nicking cleavage. We were able to improve the LOD by 100 times (Fig. 1E and 1G). Meanwhile, we also observed a high background, which makes it difficult to further distinguish targets of concentrations lower than 100 fM (Fig. 1F). To address this challenge, we further introduced a blocking DNA that competitively binds and consumes P1 (Fig. 1H). This blocking strategy was found to effectively reduce the background, allowing the detection of target DNA with 1 fM LOD (Fig. 1H and 1I).

Having established proximity CRISPR Cas12a assay for the ultrasensitive detection of nucleic acids, we next expand this assay for the amplified detection of proteins. To do so, a pair of affinity ligands that bind to the same target protein but at different epitopes were conjugated to P1 and P2 (noted as P1’ and P2’). To allow flexible binding to the target protein, we also designed a 15-nt poly-thymine spacer that separate the ligand and complementary domain (Fig. 2A). Again, the two probes do not hybridize in the absence of the target (Tm = 17 °C). However, the affinity binding to the target protein brings P1’ and P2’ into proximity, leading to the formation of a stable duplex (Fig. S2). As a result, the same CRISPR-targetable DNA barcode can be generated upon proximity primer extension, which activates Cas12a for digesting the fluorogenic substrates.

**Fig. 2.**
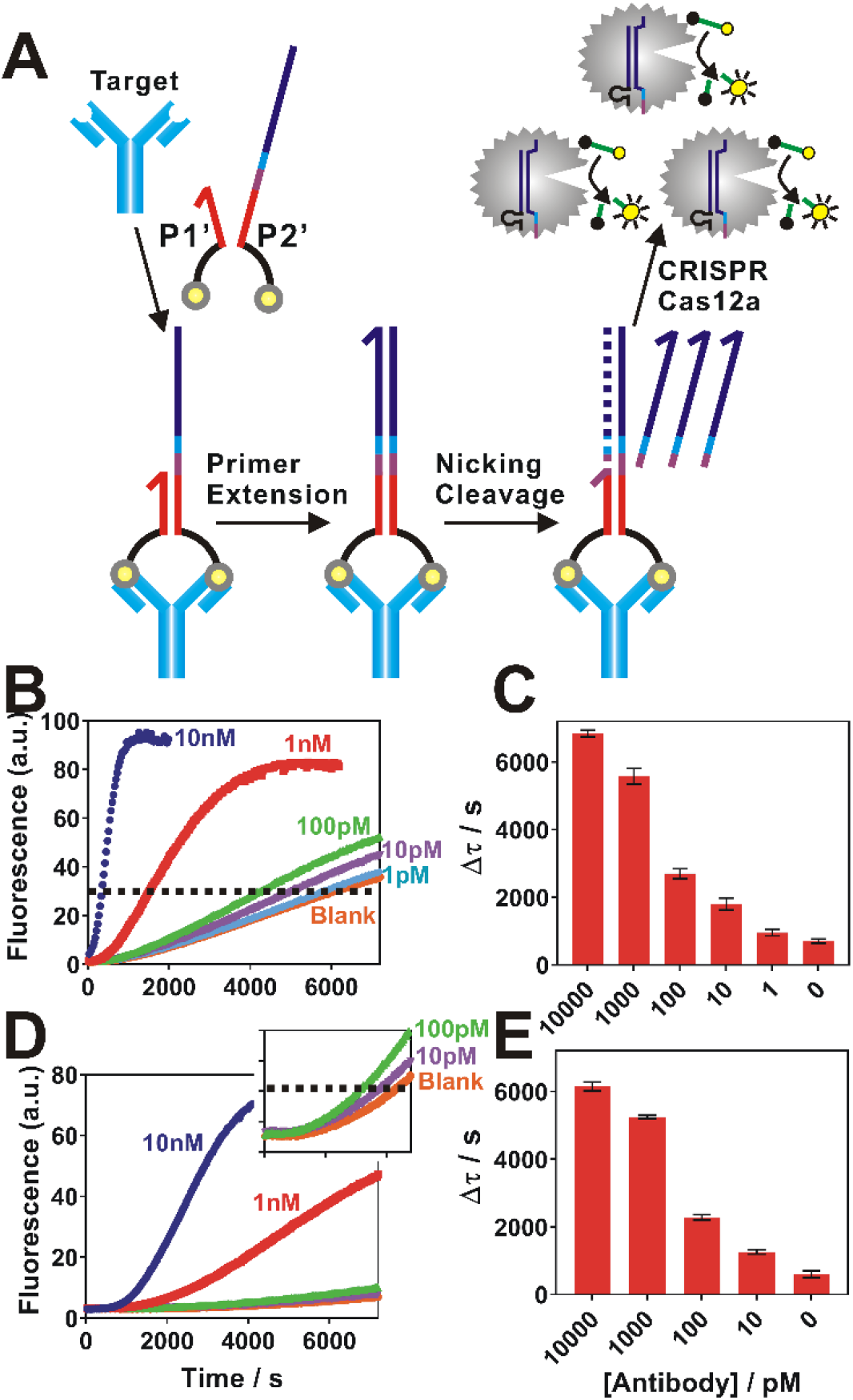
(A) Schematic illustration of the detection of antibodies using proximity CRISPR Cas12a integrated with nicking cleavage. (B, C) Detection of anti-biotin antibodies with concentrations ranging from 1 pM to 10 nM in buffer. (D, E) Detection of anti-biotin antibodies in 10-time diluted human serum samples. Each error bar represents one standard deviation from triplicate analyses.

As sensitive detection of antibodies in clinical samples plays critical roles for the diagnosis of infectious and autoimmune diseases,^7–9^ we chose anti-biotin antibody as a testbed for our assay. Correspondingly, biotin was used as the affinity ligand (Fig. 2A). In our experiments, concentrations of P1’ and P2’ and polymerase were found to be critical to maximize the binding-induced primer extension and minimize background (Fig. S3 and S4). We then integrated the assay with nicking endonuclease to further amplify the detection signal. As expected, the concentration of nicking endonuclease also significantly influences the analytical performance of the assay (Fig. S5). Under optimal experimental conditions, we were able to quantify antibodies with a LOD of 1 pM (Fig. 2B and 2C). Our assay is comparable with commonly used immunoassays in terms of sensitivity but is performed in a homogeneous solution without the need for any immobilization or washing steps. We were also able to quantify antibodies in 10-time diluted human serum samples with a LOD of 10 pM (Fig. 2D and 2E), suggesting that our assay can potentially be adapted to clinical uses.

## Conclusions

In conclusion, we have introduced a proximity CRISPR Cas12a strategy that enables Cas12a as a universal amplifier for the flexible detection of nucleic acids and proteins. DNA has long been recognized as amplifiable barcodes for the ultrasensitive detection of nucleic acids and proteins,^10–18^ with successful examples ranging from immuno-polymerase chain reactions (PCR),^11^ to proximity ligation/extension assays,^12–15^ and to biobarcode assays.^16–18^ Our proximity CRISPR assay further enriches the toolbox for ultrasensitive biomolecular detection by offering an isothermal, wash-free, and highly adaptable alternative to the existing ones. Our success in combining CRISPR detection with proximity binding also, for the first time, expands the powerful CRISPR diagnostics (CRISPR Dx) to the sensitive detection of protein-based biomarkers. Our ongoing efforts include to further improve the assay sensitivity by introducing exponential amplification components and to deploy our technology to real clinical and biological samples.

## Materials and Methods

### Materials

EnGen Lba Cas12a (Cpf1), 10 × NEBuffer™ 2.1 Buffer, Klenow Fragment (3’→5’ exo-), 10 × NEBuffer™ 2, Deoxynucleotide (dNTP) Solution Mix, nicking endonuclease (Nb.BbvCI) were purchased from New England Biolabs Ltd. (Whitby, ON, Canada). Anti-biotin antibodies were purchased from Thermo fisher Scientific (Mississauga, ON, Canada). Human serum, magnesium chloride hexahydrate (MgCl2⋅6H2O), and 100×Tris−EDTA (TE, pH 7.4) buffer were purchased from Sigma-Aldrich (Mississauga, ON, Canada). NANOpure H2O (> 18.0 MΩ), purified using an Ultrapure Mili-Q water system, was used for all experiments. All DNA samples and the guide RNAs were purchased from Integrated DNA Technologies (Coralville, IA) and purified using high-performance liquid chromatography.

### Nucleic acid detection using proximity CRISPR Cas12a assay

For a typical test, a 50 µL reaction mixture contained 5 µL of 100 nM P1, 5 µL of 100 nM P2, 10 µL of varying concentrations of genetic target, 3.3 mmol of dNTPs, 5 units of Klenow Fragment and 0.5 unit of nicking endonuclease in 1X NEBuffer™ 2. The solution was incubated at 37 °C for 20 min. A 50 µL enzyme solution which contains 30 nM of Cas12a, 30 nM of gRNA and 60 nM of signal reporter in 1× NEBuffer™ 2.1 was added. Fluorescence was measured immediately after transferring the reaction mixture to a 96-well microplate and kept measuring every 30s for 2 hours at 37 °C using a SpectraMax i3 multi-mode microplate reader (Molecular Devices) with excitation/emission at 485/515 nm.

### Proximity CRISPR Cas12a assay with blocking DNA

5 µL of 100 nM P1, 5 µL of 100 nM P2, 5 µL of 200 nM blocking DNA and 10 µL of genetic target with varying concentrations were incubated at 37 °C for 30 min. This reaction mixture was then added with 3.3 mmol of dNTPs, 5 unit of Klenow Fragment and 0.5 unit of nicking endonuclease in 1X NEBuffer™ 2 to a final volume of 50 µL. The solution was incubated at 37 °C for another 20 min. A 50 µL enzyme solution containing 30 nM of Cas12a, 30 nM of gRNA and 60 nM of signal reporter in 1× NEBuffer™ 2.1 was added. Fluorescence was measured immediately after transferring the reaction mixture to a 96-well microplate and kept measuring every 30s for 2 hours at 37 °C.

### Antibody detection using proximity CRISPR Cas12a assay

For a typical test, 5 µL of 100 nM P1’, 5 µL of 100 nM P2’ and 10 µL of varying concentrations of antibody were first mixed and incubated at 37 °C for 30 min. A 30 µL enzyme solution containing 3.3 mmol of dNTPs, 5 unit of DNA Polymerase (Klenow Fragment) and 0.5 unit of nicking endonuclease (Nb.BbvCI) in 1X NEBuffer™ 2 was then added. The solution was incubated at 37 °C for 20 min. Another 50 µL enzyme solution containing 30 nM of Cas12a, 30 nM of gRNA and 60 nM of signal reporter in 1× NEBuffer™ 2.1 was added. Fluorescence was measured immediately after transferring the reaction mixture to a 96-well microplate and kept measuring every 30s for 2 hours at 37 °C.

## End Matter

### Author Contributions and Notes

Y. Li and H. Mansour contribute equally to this work.

## Acknowledgments

We thank the Fundamental Research Funds for the Central Universities, National Sicences and Engineering Research Council of Canada, the Ontario Ministry of Research, Innovation and Science, and the Brock University Start-Up Fund for the financial support.xs

---

**Fig. S1.**
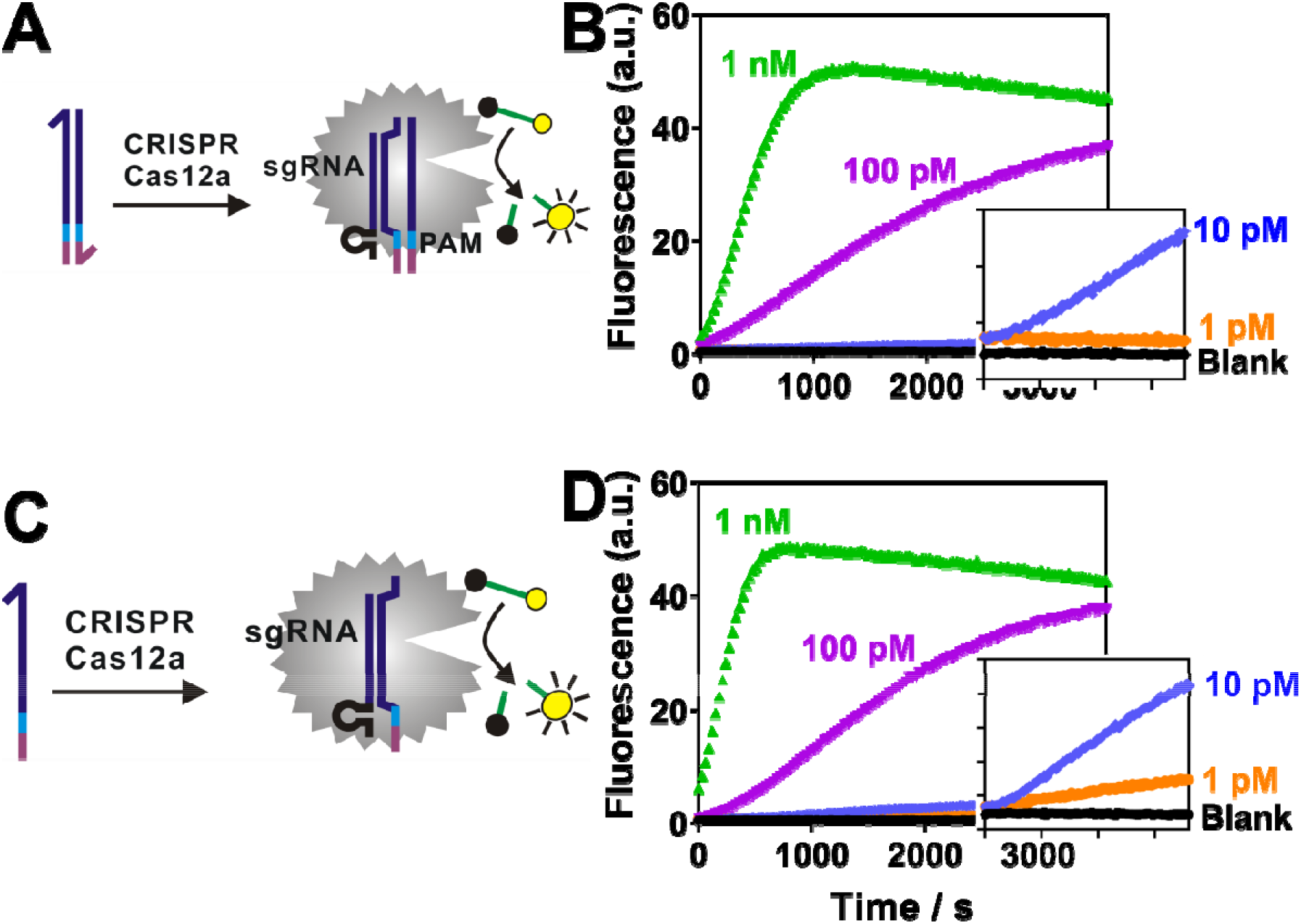
Direct detection of double-stranded DNA (dsDNA) (A, B) or single-stranded DNA (ssDNA) (C, D) using direct CRISPR RNA (crRNA) recognition and Cas12a cleavage. The limit of detection (LOD) was determined to be 10 pM for dsDNA (B) and 1 pM for ssDNA (D).

**Fig. S2.**
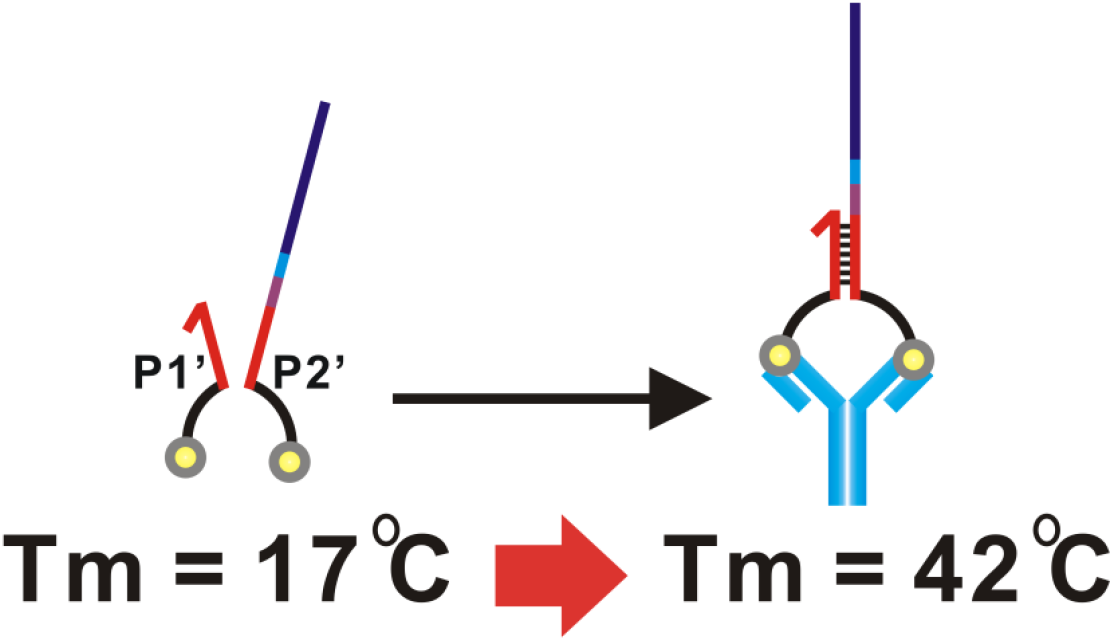
Estimated melting temperature (Tm) between P1’ and P2’ in the absence or presence of the target antibody. In absence of the target antibody, the estimated T_m_ = 17 °C by NuPack, suggesting that P1’ and P2’ do not hybridize at 37 °C. In the presence of the target, the affinity binding to the target protein brings P1’ and P2’ into proximity, leading to the formation of a stable duplex with an estimated Tm of 42 °C.

**Fig. S3.**
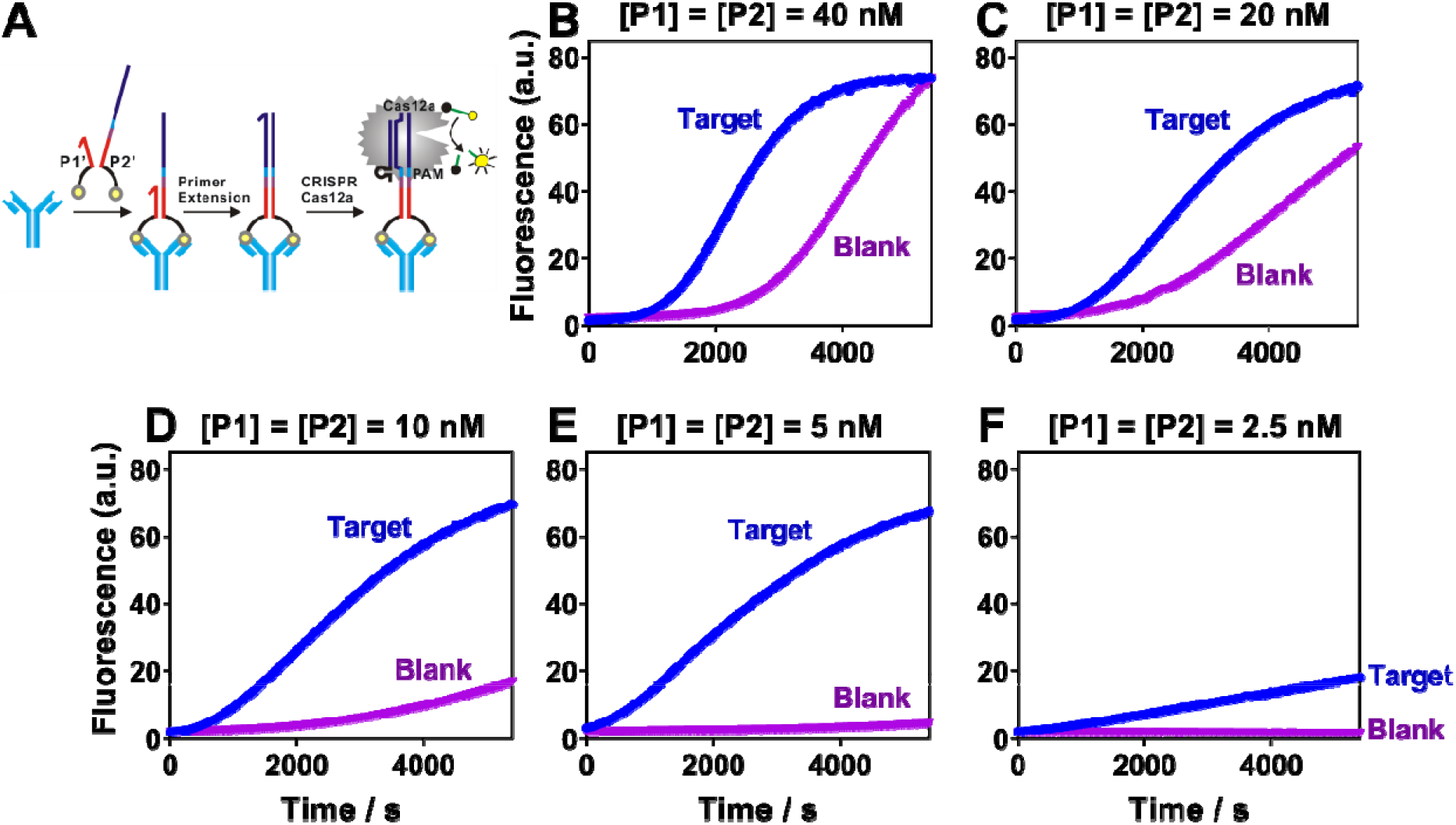
Optimization of the concentrations of proximity probes P1’ and P2’ for protein analysis. (A) Schematic illustration of the detection of protein using a binding-induced primer extension and then CRISPR Cas12a amplification. (B-F) Binding-induced primer extension using varying concentrations of P1’ and P2’ from 40 nM to 2.5 nM. The optimal concentration of P1’ and P2’ is 5 nM as it maximizes the target-dependent fluorescence signal and minimizes background signal. [Anti-biotin] = 5 nM, [Polymerase] = 1 U.

**Fig. S4.**
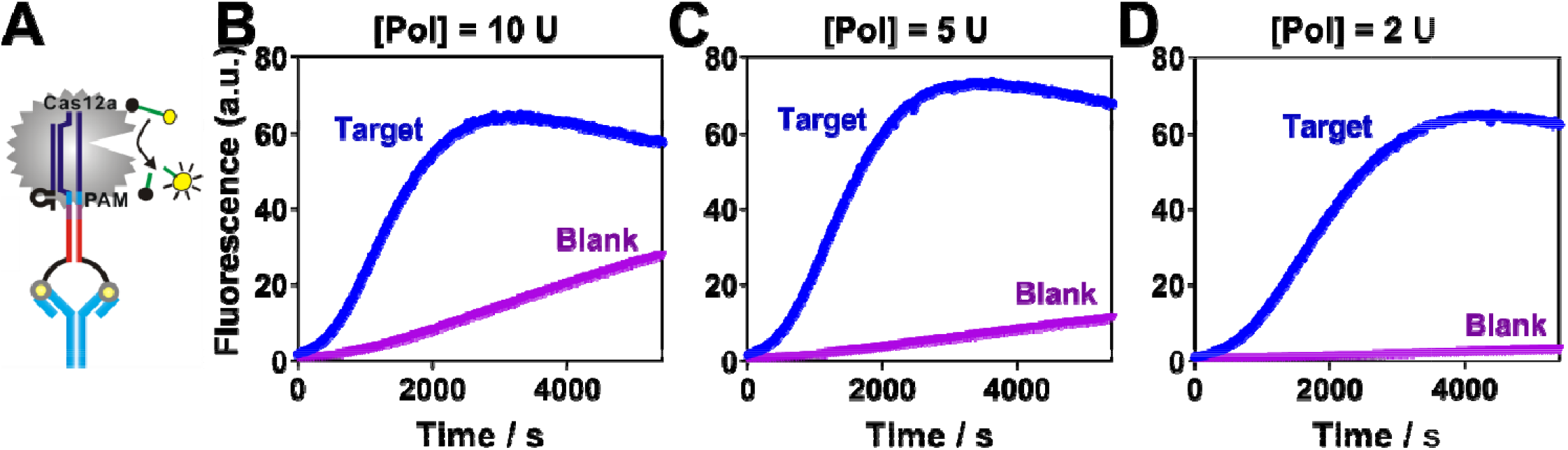
Optimization of the concentrations of DNA polymerase (Klenow Fragment, unit) for protein analysis. (A) The detection of protein was achieved using a binding-induced primer extension and then CRISPR Cas12a ampification. (B-D) Detection of anti-biotin antibody using varying concentrations of DNA polymerase. The optimal amount of Klenow Fragment was found to be 5 units, as it maximizes detection signals and kinetics while maintains a reasonably low background. [Anti-biotin] = 5 nM, [P1’] = [P2’] = 5 nM.

**Fig. S5.**
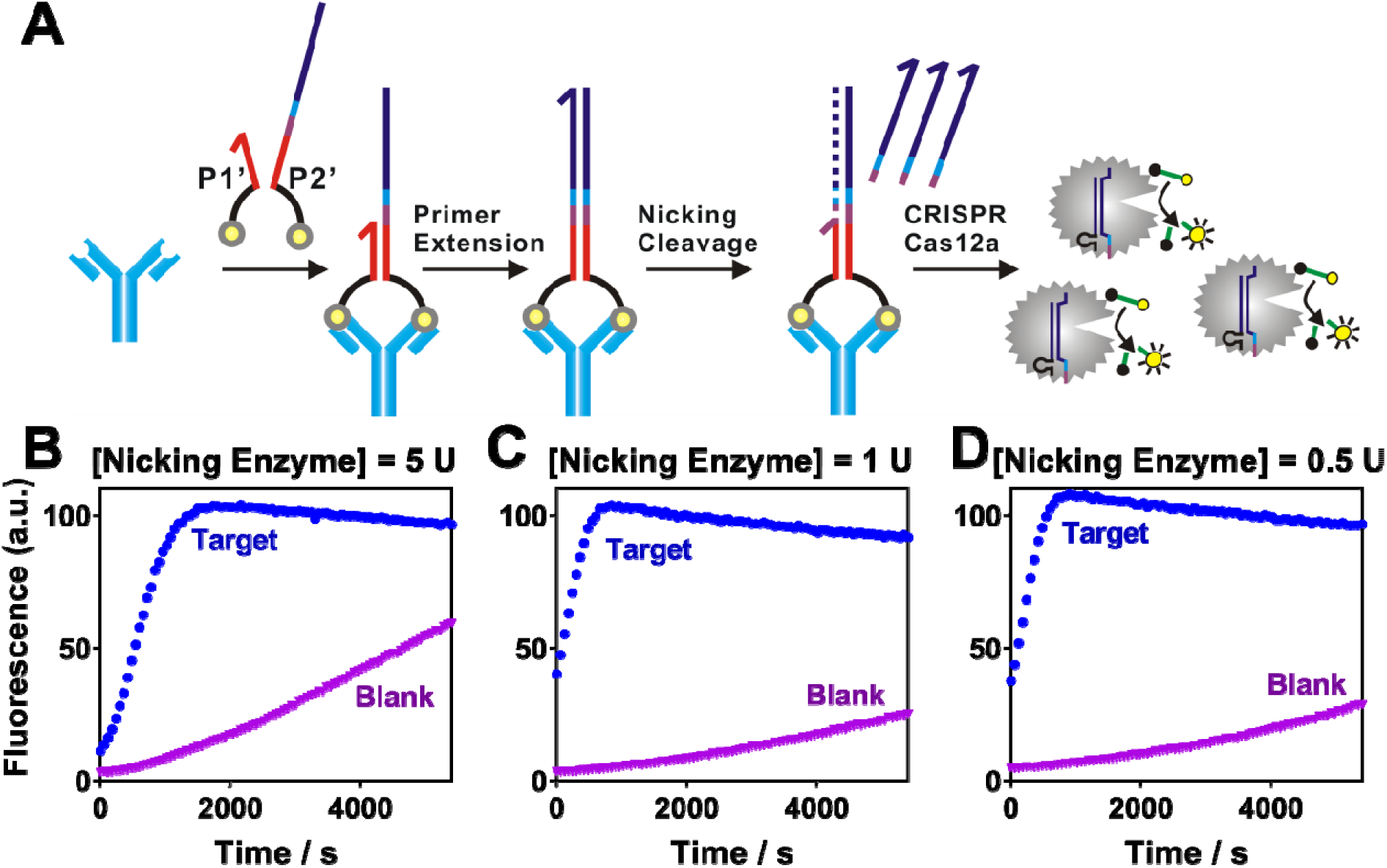
Optimization of nicking endonuclease for the proximity CRISPR Cas12a assay. (A) Schematic illustration of antibody detection using priximity CRISPR Cas12a combined with nicking cleavage. (B-D) Detection of anti-biotin antibody using varying concentrations of nicking endonuclease from 0.5 U to 5 U. The optimal amount of nicking endonuclease was found to be 0.5 units, as it maximizes detection signals and minimizes the background. [Anti-biotin] = 5 nM, [P1’] = [P2’] = 5 nM, [Polymerase] = 5 U.

