## Supplementary material for "A Universal Proximity CRISPR Cas12a Assay for Ultrasensitive Detection of Nucleic Acids and Proteins": Supplmental material

**1. Materials.** EnGen Lba Cas12a (Cpf1), 10 × NEBuffer™ 2.1 Buffer, Klenow Fragment (3'→5' exo-), 10 × NEBuffer™ 2, Deoxynucleotide (dNTP) Solution Mix, nicking endonuclease (Nb.BbvCI) were purchased from New England Biolabs Ltd. (Whitby, ON, Canada). Anti-biotin antibodies were purchased from Thermo fisher Scientific (Mississauga, ON, Canada). Human serum, magnesium chloride hexahydrate ($MgCl_2 \cdot 6H_2O$), and 100×Tris−EDTA (TE, pH 7.4) buffer were purchased from Sigma-Aldrich (Mississauga, ON, Canada). NANOpure $H_2O$ (> 18.0 MΩ), purified using an Ultrapure Mili-Q water system, was used for all experiments. All DNA samples and the guide RNAs were purchased from Integrated DNA Technologies (Coralville, IA) and purified using high-performance liquid chromatography. The DNA and RNA sequences are outlined in Table S1.

**Table S1. DNA sequences and modifications.**

| Name | Sequence (5'→3') |
|---|---|
| **Nucleic Acid Detection** | |
| P2 (Template) | GCT TGT GGC CG TTTA CGT CGC CGT CCA GCT CGA CCTCAGC CGTAGA TT GAC TCT GGC TTT-InvT |
| P1 (Primer) | ATC TCT CTG AAG TT TCTACG |
| Blocking DNA | TTT TTT CGTAGA[c] |
| Target | AAA AGA TAA CAA GAA AGAC AAA GCC AGA GTC CTT CAG AGA GA TAC AGA AAC TCT AAT TCA |
| **Protein Detection** | |
| P2' (Template) | GCT TGT GGC CG TTTA CGT CGC CGT CCA GCT CGA CCTCAGC ATGCGTAGA TTT TTT TTT TTT TTT-Biotin |
| P1' (Primer) | Biotin-TTT TTT TTT TTT TTT TCTACG |
| **CRISPR-Cas12a** | |
| crRNA | UAA UUU CUA CUA AGU GUA GAU CGU CGC CGU CCA GCU CGA CC |
| Signal Reporter | FAM-TTATT-Quencher |

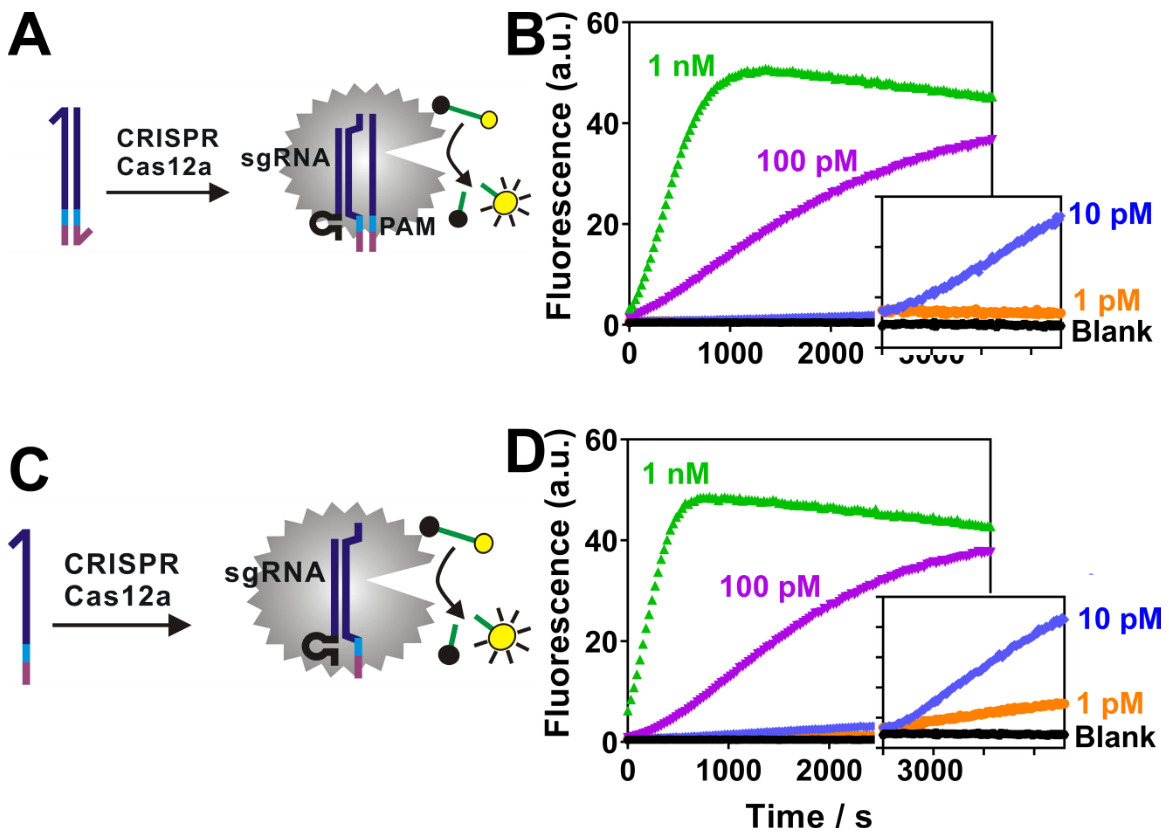


**Figure S1**. Direct detection of double-stranded DNA (dsDNA) (**A**, **B**) or single-stranded DNA (ssDNA) (**C**, **D**) using direct CRISPR RNA (crRNA) recognition and Cas12a cleavage. The limit of detection (LOD) was determined to be 10 pM for dsDNA (**B**) and 1 pM for ssDNA (**D**).

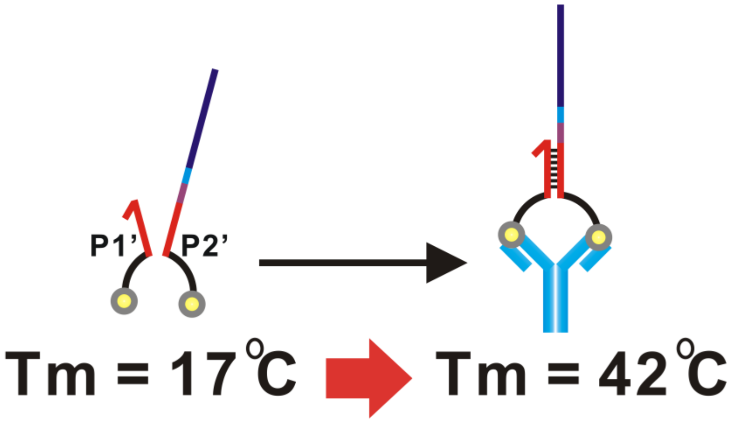


**Figure S2.** Estimated melting temperature ($T_m$) between P1' and P2' in the absence or presence of the target antibody. In absence of the target antibody, the estimated $T_m$ = 17 °C by NuPack, suggesting that P1' and P2' do not hybridize at 37 °C. In the presence of the target, the affinity binding to the target protein brings P1' and P2' into proximity, leading to the formation of a stable duplex with an estimated $T_m$ of 42 °C.

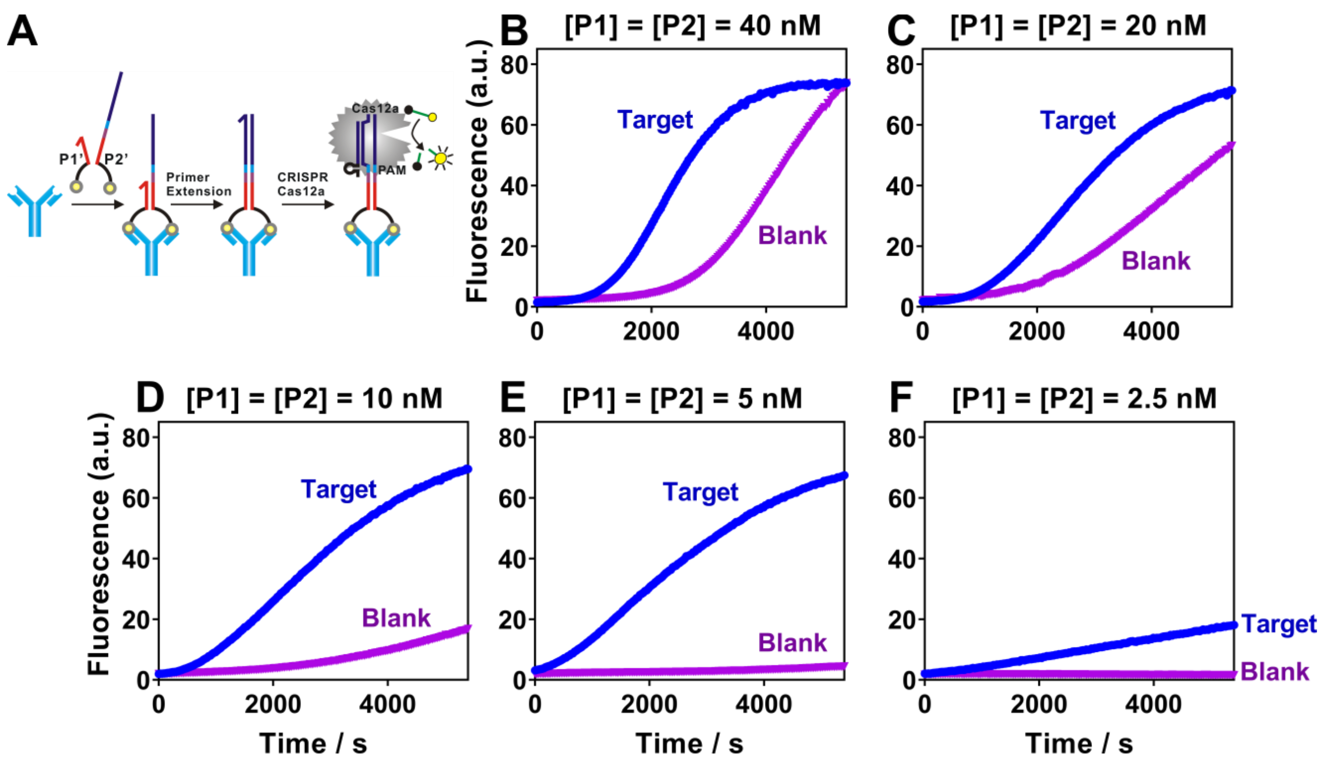


**Figure S3**. Optimization of the concentrations of proximity probes P1' and P2' for protein analysis. (**A**) Schematic illustration of the detection of protein using a binding-induced primer extension and then CRISPR Cas12a amplification. (**B-F**) Binding-induced primer extension using varying concentrations of P1' and P2' from 40 nM to 2.5 nM. The optimal concentration of P1' and P2' is 5 nM as it maximizes the target-dependent fluorescence signal and minimizes background signal. [Anti-biotin] = 5 nM, [Polymerase] = 1 U.

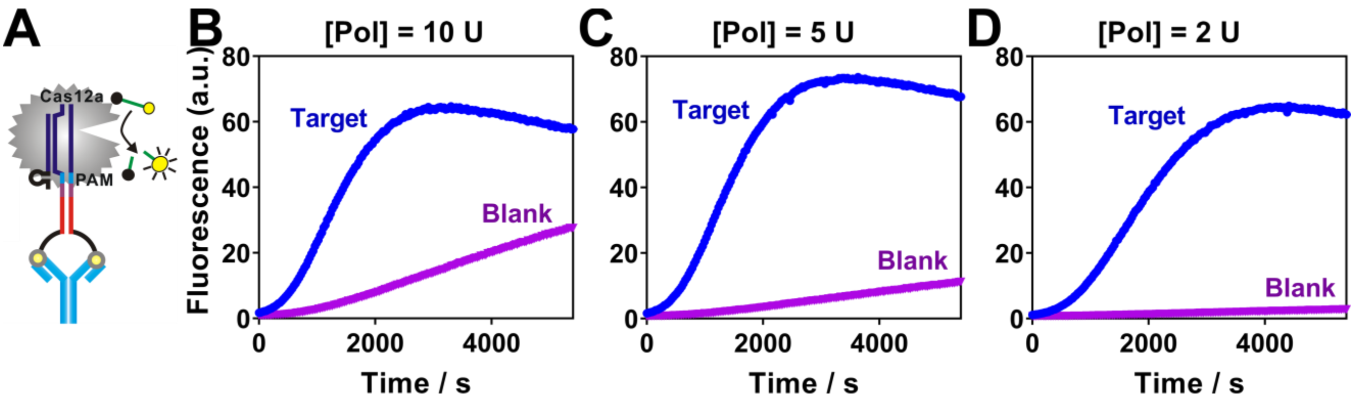


**Figure S4**. Optimization of the concentrations of DNA polymerase (Klenow Fragment, unit) for protein analysis. (**A**) The detection of protein was achieved using a binding-induced primer extension and then CRISPR Cas12a ampification. (**B-D**) Detection of anti-biotin antibody using varying concentrations of DNA polymerase. The optimal amount of Klenow Fragment was found to be 5 units, as it maximizes detection signals and kinetics while maintains a reasonably low background. [Anti-biotin] = 5 nM, [P1'] = [P2'] = 5 nM.

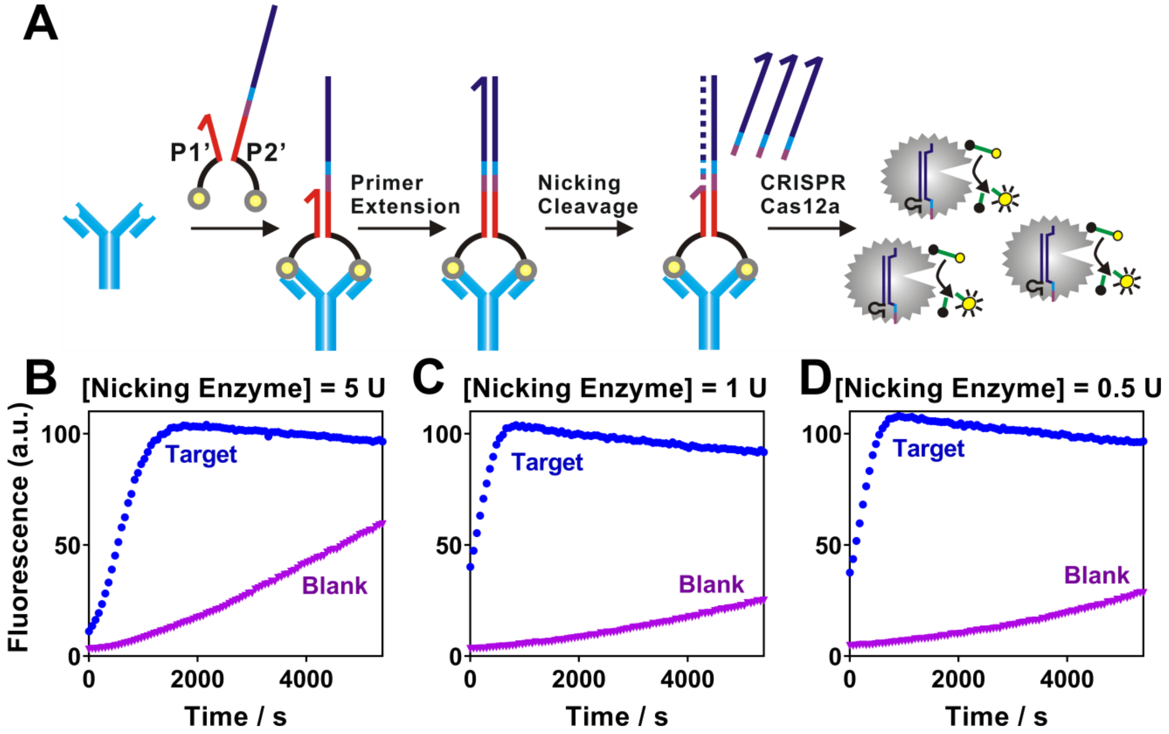


**Figure S5.** Optimization of nicking endonuclease for the proximity CRISPR Cas12a assay. (**A**) Schematic illustration of antibody detection using priximity CRISPR Cas12a combined with nicking cleavage. (**B-D**) Detection of anti-biotin antibody using varying concentrations of nicking endonuclease from 0.5 U to 5 U. The optimal amount of nicking endonuclease was found to be 0.5 units, as it maximizes detection signals and minimizes the background. [Anti-biotin] = 5 nM, [P1'] = [P2'] = 5 nM, [Polymerase] = 5 U.
